# Sleep spindle dynamics suggest over-consolidation in post-traumatic stress disorder

**DOI:** 10.1101/2021.07.29.453342

**Authors:** A.C. van der Heijden, W.F. Hofman, M. de Boer, M.J. Nijdam, H. J. F. van Marle, R. A. Jongedijk, M. Olff, L.M. Talamini

**Author notes:** Corresponding author: A.C. van der Heijden (Christa).

## Abstract

Devastating and persisting traumatic memories are a central symptom of post-traumatic stress disorder (PTSD). Sleep problems are highly co-occurrent with PTSD and intertwined with its etiology. Notably, sleep hosts memory consolidation processes, supported by sleep spindles (11-16 Hz). Here we assess the hypothesis that intrusive memory symptoms in PTSD may arise from excessive memory consolidation, reflected in exaggerated spindling. We use a newly developed spindle detection method, entailing minimal assumptions regarding spindle phenotype, to assess spindle activity in PTSD patients and traumatized controls (n=2×14, matched on gender). Our results show increased spindle activity in PTSD, which positively correlates with daytime intrusive memory symptoms. Together, these findings provide a putative mechanism through which profound sleep disturbance in PTSD may contribute to memory problems. Due to its uniform and unbiased approach, the new, minimal assumption spindle detection method seems a promising tool to detect aberrant spindling in psychiatric disorders.

## Introduction

The majority of individuals (over 70%) experience a traumatic event in their lifetime, leading to a lifetime prevalence of post-traumatic stress disorder (PTSD) in about 7% (1,2) A characteristic symptom of PTSD is involuntary, distressing recall of the traumatic event, which can occur in the form of intrusive memories, flashbacks or nightmares. Besides these memory disturbances, PTSD patients suffer from severe sleep problems (3), like insomnia, nightmares, distressed awakenings, nocturnal panic attacks, and sleep terrors (4). These sleep problems, which occur in 70-90% of patients (5) and often predate the trauma (6), are thought to play an important role in PTSD genesis and maintenance (5,7,8). Accordingly, sleep problems, either predating or following trauma, are a strong predictor for the development of PTSD (9,10) They are also a frequent residual symptom after successful psychopathology-oriented treatment (11) and then constitute a strong predictor for relapse.

Interestingly, sleep importantly supports memory consolidation (12) through sleep spindles (13–16): waxing and waning oscillations in the sigma frequency band (11-16 Hz), generated by thalamocortical feedback loops. Sleep spindles have been shown to reflect the reprocessing of individual memory traces (13) and are causally related to emotional memory consolidation (16). A recent study in our lab demonstrated an increase in relative sigma power in patients with PTSD compared to healthy subjects, peaking in right-frontal areas (17), which are involved in the processing of emotional and self-relevant memories (18). As sigma power is highly correlated to sleep spindle activity (19) this may reflect exaggerated spindle activity related to the reprocessing of trauma memories, which consequently might lead to intrusive memory symptoms.

A recent study that directly examined sleep spindles in PTSD (Wang 2020) did not report increased spindle activity, although heightened slow-spindle oscillatory frequency over frontal brain regions was observed. However, we will argue hereafter that the methods used to detect spindles in this study, although commonly used, have drawbacks that limit their suitability to assess spindle abnormalities associated to, for instance, psychological disorders.

A pitfall in most previous spindle studies is the use of non-uniform,*a priori* criteria for the definition of sleep spindles. These criteria lack evidence of physiological delineation and often include high amplitude and duration thresholds, leading to arbitrary cut-offs in the analyzed spindle activity. Spindle detection methods of this kind have several important drawbacks: for one, notable parts of spindle activity are not considered in the analyses. Second, arbitrary choices with respect to the aforementioned criteria and their calculation can lead to large differences in results, and thus in difficulties comparing studies.

To analyze sleep spindles in an unbiased manner, a new, minimal assumptions, spindle analysis method was developed. This method defines spindle events in terms of waxing and waning dynamics in the sigma band. It detects discrete spindle events of virtually any size and allows examination of the morphology of each event. A minimal assumption method may be particularly relevant to spindle analyses in mental disorders, since standard assumptions may not apply and might even hamper the detection of deviant physiology representative of the disorder.

In order to investigate this hypothesis our first aim was to reliably quantify spindle activity in PTSD. Specifically, we examined sleep spindle activity, including several spindle event parameters (e.g., amplitude and duration), in police officers and war veterans with and without PTSD. We, furthermore, assessed whether changes in spindle activity in patients with PTSD were associated to intrusive memory complaints. To understand how potential changes at the level of discrete spindle events relate to spectral EEG measures absolute sigma power was also assessed.

## Methods & Materials

### Participants

The participants consisted of patients with chronic PTSD (n=14), according to the Clinician-Administered PTSD Scale (CAPS) for DSM IV (20), and trauma-exposed controls (n=14). Trauma-exposed controls had experienced a traumatic event (as defined by DSM-IV), but did not meet de diagnostic criteria for PTSD. Participants with PTSD were recruited at ARQ Centrum’45, a Dutch national center for the diagnosis and treatment of PTSDTrauma-exposed controls were recruited through police departments, and veterans’ centers. The two groups were matched on age, gender, profession, and educational level. No differences in alcohol-related disorders and use were found between groups. Participants were asked to refrain from medication use prior to the experiment; however, for six of the PTSD patients, usage of psychotropic medication could not be interrupted (five patients were on serotonin reuptake inhibitors (Paroxetine, Venlafaxine, Sertraline, Citalopram), whereas one patient used a benzodiazepine (Temazepam). Subjects were excluded in case of acute suicidality, presence of a psychotic or bipolar disorder, depression with psychotic features, excessive substance related disorder over the past 3 months before inclusion, history of neurological or sleep disorders (prior to PTSD onset), a habitual sleep pattern with less than 6 hours sleep per night, or a sleep window outside 10 PM till 10 AM. The study protocol was approved by the Medical Ethical Committee of the Amsterdam Medical Center (AMC). All participants gave written informed consent.

### Polysomnography

For polysomnographic recordings, subjects were given the opportunity to sleep undisturbed for nine hours during a lights-off period, starting between 11 and 12 PM, depending on habitual sleep times. Polysomnography, recorded in the context of a more extensive program on PTSD and sleep (17), was performed using ambulatory 16-channel Porti amplifiers (TMS-i) and Galaxy sleep analysis software (PHI-international). Polysomnography included an EEG recording (F3, F4, C4, O2, referenced to the average of the mastoids (M1, M2), two EOG electrodes monitoring eye-movements, and two electrodes for submental EMG (for the full montage see (17)). All signals were sampled at a rate of 512Hz. Sleep stages were scored visually, according to the standard AASM criteria (21).

### Sigma power

The sigma frequency content of the EEG (11-16 Hz) was analyzed using fast Fourier transform-based spectral analysis (4s time windows with 50% overlap, 0.25Hz bin size; Hamming window) on right-frontal electrode, F4, for N2 sleep. First, through visual inspection of the data, EEG epochs containing artifacts were removed. Next, for each frequency bin in the sigma band, the power per 30s epoch was computed and summated over all epochs. Finally, absolute power bins were merged across frequencies, and power was divided by time in N2 for each subject.

### Minimal assumptions spindle analysis

Automatized detection of discrete spindle events and computation of spindle event parameters, used a minimal assumptions spindle analysis algorithm (module within Galaxy sleep analysis software), provided by PHI international, Amsterdam, The Netherlands. Specifically, sleep EEG data of electrode F4 (referenced to the average of the mastoids), was filtered in the sigma range (11-16 Hz) using a Finite Impulse Response (FIR) filter built into the algorithm. With a moving window of 0.2 seconds, the standard deviation of the filtered signal was calculated, resulting in a spindle envelope with a sample rate of 512 Hz. The envelope represents the burst-like shape of the signal. To convert the standard deviation of the filtered signal to μV, the standard deviation was multiplied by √2. To define peaks and troughs, the first derivative (slope) of the envelope was calculated, after first down-sampling the data to 5 Hz (achieved by taking the root mean square of the envelope data over sequential stretches of 0.2 seconds). A peak was defined if the slope changed from positive to negative, a trough vice versa. A hysteresis of 0.2 μV was used to neglect low amplitude background noise in the envelope, while still including a broad range of spindle events for analyses. All spindle events with amplitude between 5 and 35 μ V were analyzed.

For each spindle event four parameters were calculated: duration, peak amplitude (absolute maximal amplitude), waxing amplitude (difference between the absolute level of the peak and the absolute level of the trough prior to the peak), and waning amplitude (difference between the absolute level of the peak and the absolute level of the trough after the peak) (figure 1). As this method detects spindle events in a wide range of sizes, it delivers rich, comprehensive data about spindle activity while offering many possibilities for analysis.

**Figure 1.**
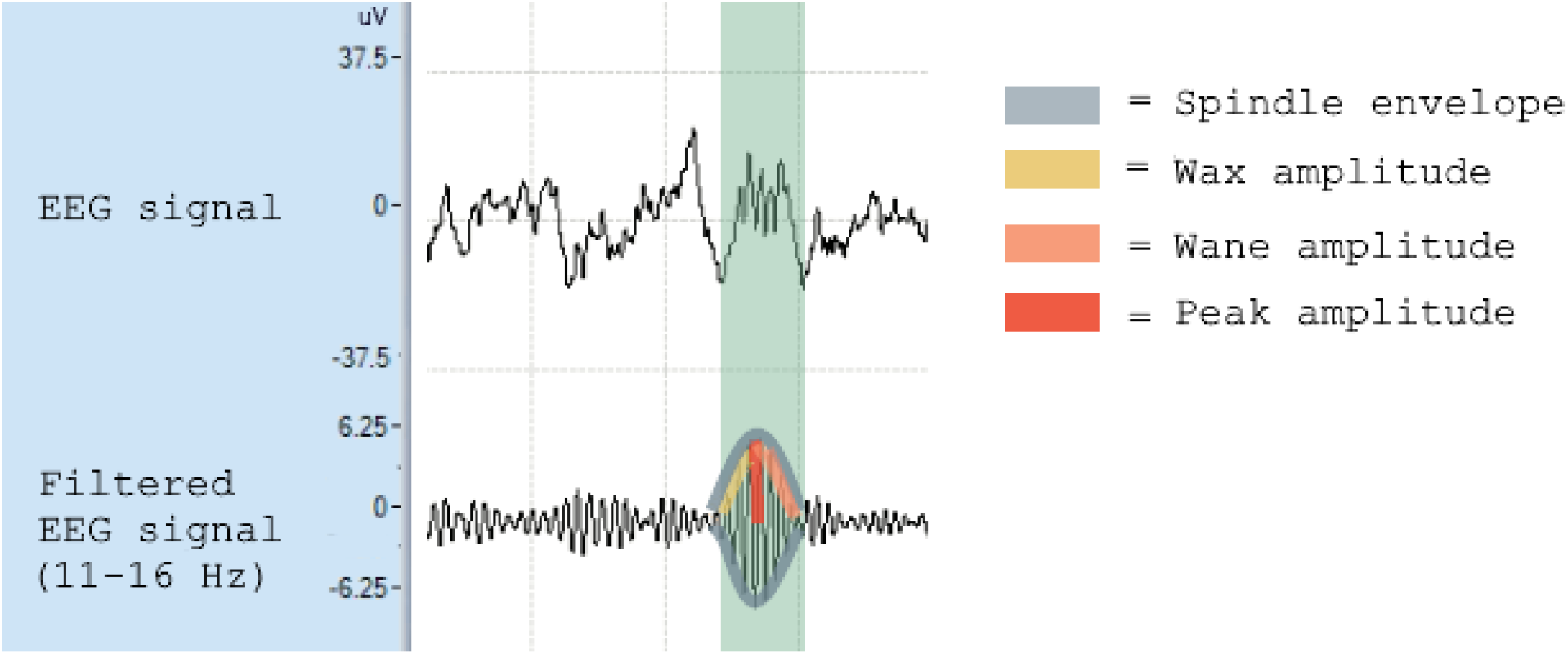
Spindle detection method and definition of spindle parameters The EEG signal was filtered into spindle frequency (11-16 Hz). With a moving window, the standard deviation of the filtered signal was calculated, resulting in a spindle envelope that follows the burst-like shape of the signal. From the spindle envelope, the following amplitude parameters are calculated: peak amplitude (absolute maximal amplitude), waxing amplitude (difference between the absolute level of the peak and the absolute level of the trough prior to the peak), and waning amplitude (difference between the absolute level of the peak and the absolute level of the trough after the peak)

### Spindle activity index

In order to correlate spindle activity to clinical measures reflecting memory intrusions, we developed an index that captured the deviations of the PTSD spindle envelope into a single measure. The correlation matrix of the spindle parameters that were changed in the PTSD group (peak, wax and wane amplitude) showed that wax and wane amplitude were highly intercorrelated (r=0.989, p<0.01). Thus, these two variables have similar information content. Wax and wane amplitude were each less correlated with peak amplitude (wax amplitude vs peak amplitude: r=−0.651, p<0.01; wane amplitude vs peak amplitude: r=−0.660, p<0.01). Therefore, a Spindle Activity Index (SAI) was calculated as peak amplitude divided by the average of the wax- and wane amplitude.

### Statistical analyses

As most of the data was not normally distributed, statistical analyses were performed using non-parametric methods and all averages in the results are given as mean ranks.

## Results

### Sigma power

Absolute right-frontal sigma power in N2 was compared between PTSD and controls using a two-tailed Mann-Whitney test. Right-frontal sigma power was increased in PTSD patients relative to controls by 144% (ptsd=17.36, ctrl=10.38, u=44, p=0.02).To control for possible medication effects on right-frontal sigma power, a Quade’s ANCOVA was conducted in which medication intake (yes/no) was added as a covariate. In this analysis, the increase in right-frontal sigma power in PTSD patients remained significant (F(1,27)=4.323, p=0.048), suggesting a PTSD-related sigma power increase independent of medication effects.

### Spindle parameters

Averages for each spindle parameter were calculated over the entire distribution and compared between PTSD patients and controls, using a Mann-Whitney U test. These tests revealed higher mean peak amplitude in PTSD subjects (=17.93) compared to controls (=11.07; U=50, p=0.027). In contrast, local fluctuations between peaks and troughs in the spindle envelope were shallower in PTSD relative to controls (waxing amplitude: ptsd=10.14, ctrl=18.86, U=37, p=0.005; waning amplitude: ptsd=10.14, ctrl=18.86, U=37, p=0.004). No statistically significant difference in the duration of these local envelope fluctuations was found (PTSD=12.00, ctrl=17.00, p=0.108). The combined evidence points to an inflated spindle envelope in PTSD patients, which remains at high amplitude levels, rarely dropping to low levels. This, in turn, suggests increased spindle activity in PTSD compared to controls.

To control for medication effects on these spindle parameters, Quade’s ANCOVAs were conducted, with medication intake (yes/no) as a covariate. For peak amplitude, the differences between groups reduced to trend-level significance (F(1,28)=3.233, p=0.084) after controlling for medication intake. As such, an influence of medication on these results cannot be entirely excluded. Differences between groups for local trough to peak fluctuations were unaffected by medication effects (F(1,28)=6,940 p=0.014). In summary, while medication effects on the spindle envelope parameters cannot be excluded, they did not explain the differences between groups entirely.

Besides, we examined, for each spindle parameter, whether the density distribution was similar between the PTSD patient and control group, using a two-sample Kolmogorov-Smirnov test. This assessed whether differences between groups might be specific to a particular part of the distribution (e.g. to the smallest or the largest spindles). No differences were found (all p’s>0.1, see Supplemental information S1).

### Spindle activity & intrusive memory symptoms

To analyze whether changes in spindle activity were associated with PTSD intrusive memory complaints, spindle activity, indexed by the SAI (see methods) was correlated to the intrusive memory symptom score of the diagnostic tool for PTSD (total score of items under criterion B in CAPS-IV) using Spearman’s rank correlation coefficient test. This revealed a positive correlation (r_s_=0.383, p=0.045), indicating a link between increased spindle activity and memory intrusions of the traumatic event.

Next, we explored whether this correlation was present for all types of intrusive memory symptoms, by correlating the SAI to individual scores on the B items of the CAPS-IV. SAI was significantly correlated to memory intrusions (CAPS-B1 r_s_=0.477, p=0.010), re-experiencing (flashbacks of) the traumatic event (CAPS-B3 r_s_=0.406, p=0.032), emotional and physical responses to reminders of the trauma (respectively, CAPS-B4 r_s_=0.412, p=0.029, CAPS-B5 r_s_=0.419, p=0.026). The SAI showed only trend-level correlation to the frequency of trauma-related nightmares (CAPS-B2 r_s_=0.322, p=0.085), and no relation at all to nightmare intensity (CAPS-B2I, p=0.251, CAPS-B2, p=0.163,). Together, these findings demonstrate that increased spindle activity is predominantly linked to intrusive memory symptoms during wake, showing little or no association to nightmare-related memory intrusions.

## Discussion

Using a minimal assumption spindle detection method, this study revealed increased right-frontal spindle activity in PTSD patients compared to trauma-exposed controls. The increase was reflected in inflation of the spindle envelope and augmented absolute sigma power. Furthermore, inflation of the spindle envelope was associated with more daytime, intrusive trauma memories.

A large body of evidence supports the relation between spindle activity and memory performance (22–26). According to this literature, spindles reflect the reactivation and consolidation of cortical memory traces, enhancing post-sleep memory for the pertaining information. Spindles are largely local phenomena (27), reflecting reprocessing of the type of information encoded in a particular cortical area (28). As the right prefrontal cortex is involved in the processing of emotional and self-relevant information across sleep-wake states (18), the observed increase in spindle activity in this area may reflect the excessive reprocessing and strengthening of trauma memories during sleep. Supporting this notion, right-frontal spindle activity was positively correlated with daytime intrusive memory symptoms in PTSD patients. The lack of a similar correlation with nightmare-related trauma-memory intrusions might be related to nightmares mostly occurring during REM sleep, while spindles occur during NREM sleep. These findings suggest that PTSD-related abnormalities in NREM and REM sleep are to some extent disassociated and may have a different relation to symptomatology.

Besides sleep spindles, sleep slow oscillations (SO’s) (0.5-1.5 Hz) are important for memory consolidation (29,30). We have shown previously that SO dynamics are strongly impaired in PTSD (17). This may affect emotional memory processing in several ways: first, SO’s synchronize sleep spindles and higher frequency activity within and across cortical regions (31).This is thought to support long-range communication across the brain, including the hippocampo-cortical cross-talk underlying system-level consolidation of episodic memories (29,31). The lack of coordinating background SO activity ((in PTSD)) may undermine these ((system-level consolidation)) processes. As a result, event memory traces might be strengthened, in their original hippocampus-dependent form, with preservation of episodic detail and emotional tone, rather than being integrated into memory networks as more general memories.

Second, SO activity and deep sleep support a host of growth and recovery processes that are essential to physical and mental integrity (32,33). These processes, which include synaptic equilibration, neural waste clearance and metabolic anabolic reactions, likely contribute to sleep’s crucial role in emotional housekeeping (34). Accordingly, slow wave sleep, has been shown to support the emotional attenuation of memories in healthy subjects (35), while deep sleep deficits were correlated to failing affective attenuation in youth with PTSD (36)

Thus, taking into account our previous findings on aberrant SO dynamics in PTSD together with the current finding of increased right-frontal spindle activity, we suggest that intrusive memory symptoms in PTSD may arise from memories being (overly) strengthened in their original configuration, with rich episodic detail and high emotional tone. At the same time, disruption of general recovery processes would further compromise neural processing and, therewith, cognitive and emotional functioning.

It might here be noted that the alterations in sleep spindles and slow oscillations discussed above are likely interlinked to other changes in PTSD sleep physiology, including changes in REM sleep’s spectral topology (13), neuroendocrine (27) and autonomic regulation (37). These changes, which to some extent may reflect a state of hyperarousal during sleep, are all likely to affect information processing, as well as the normal play out of general recovery processes, during sleep.

A particular strength of our study was the use of a novel spindle detection method, involving minimal assumptions regarding spindle morphology. This method holds several benefits over traditional spindle detection methods using high, arbitrary thresholds. For one, criteria that base themselves on the norm, by definition neglect phenomenology outside of the norm, thus underestimating or altogether failing to detect abnormalities in non-norm groups or conditions. Compounding the problem, arbitrary threshold and baseline choices in traditional spindle detection methods can vastly affect analysis results, undermining findings’ robustness and comparability between studies. The minimal assumption method could increase sensitivity, reproducibility and comparability in spindle research, while combating arbitrarily chosen spindle detection threshold parameters, as urged by various experts in the field (19,24).

Our study also has limitations, including a limited sample size. While the tight matching of patient and trauma-controls on a broad range of sociodemographic variables aids the power to detect PTSD related differences, the tightly defined participant sample (treatment-seeking, mostly male, police officers and veterans, with severe, chronic PTSD) warrants caution in extrapolating to other PTSD populations.

In summary, increased right-frontal sleep spindle activity may provide a mechanism through which the profound sleep disturbances in PTSD contribute to intrusive memory symptoms. Besides their fundamental interest, these findings emphasize the need to take sleep-related symptoms into account in PTSD treatment strategies, both as a treatment target and as an outcome measure. While it might be tempting to consider treatments targeted at sleep spindle disruption, we would rather advocate treatments aimed at restoring healthy sleep, including restoration of sleep continuity and sleep depth and, therewith, normalize spindling as well as other sleep physiological parameters.

The minimal assumption spindle detection may be a powerful tool in psychiatric research due to its broad, uniform detection range, which is permissive towards pathology-associated physiological aberrancies, while increasing comparability between studies.

## Acknowledgements

We thank the staff of ARQ Centrum’45 for their involvement in patient recruitment and varied practical support that has been crucial to the realization of this study. We thank Anand Kumar from PHI International for his contribution and help regarding the spindle detection in this project.

## References

1. Kessler, Ronald C, Sonnega, A., Bromet, E.; Hughes M., Nelson CB (1995): Posttraumatic Stress Disorder in the National Comorbidity Survey. Gen Hosp Psychiatry 52: 1048–1060.

2. De Vries GJ, Olff M (2007): The Lifetime Prevalence of Traumatic Events and Posttraumatic Stress Disorder in the Netherlands. 20: 251–262.

3. Lewis C, Lewis K, Kitchiner N, Isaac S, Jones I, Bisson JI (2020): Sleep disturbance in post-traumatic stress disorder (PTSD): a systematic review and meta-analysis of actigraphy studies. Eur J Psychotraumatol 11. https://doi.org/10.1080/20008198.2020.1767349

4. Germain A (2013): Sleep disturbances as the hallmark of PTSD: Where are we now? Am J Psychiatry 170: 372–382.

5. Spoormaker VI, Montgomery P (2008): Disturbed sleep in post-traumatic stress disorder: Secondary symptom or core feature? Sleep Med Rev 12: 169–184.

6. Babson KA, Feldner MT (2010): Temporal relations between sleep problems and both traumatic event exposure and PTSD: a critical review of the empirical literature, 2009/09/01. J Anxiety Disord 24: 1–15.

7. Pace-Schott EF, Germain A, Milad MR (2015): Sleep and REM sleep disturbance in the pathophysiology of PTSD: The role of extinction memory. Biol Mood Anxiety Disord 5: 1−19.

8. Sopp MR, Brueckner AH, Schäfer SK, Lass-Hennemann J, Michael T (2019): Differential effects of sleep on explicit and implicit memory for potential trauma reminders: findings from an analogue study. Eur J Psychotraumatol 10. https://doi.org/10.1080/20008198.2019.1644128

9. Koren D, Arnon I, Lavie P, Klein E (2002): Sleep complaints as early predictors of posttraumatic stress disorder: a 1-year prospective study of injured survivors of motor vehicle accidents. Am J Psychiatry 159: 855–857.

10. Bryant RA, Creamer M, O’Donnell M, Silove D, McFarlane AC (2010): Sleep disturbance immediately prior to trauma predicts subsequent psychiatric disorder. Sleep 33: 69–74.

11. Smid GE, van der Meer CAI, Olff M, Nijdam MJ (2018): Predictors of Outcome and Residual Symptoms Following Trauma-Focused Psychotherapy in Police Officers With Posttraumatic Stress Disorder. J Trauma Stress 31: 764–774.

12. Stickgold R (2005): Sleep-dependent memory consolidation. Nature 437: 1272–1278.

13. Cox R, Hofman WF, de Boer M, Talamini LM (2014): Local sleep spindle modulations in relation to specific memory cues. Neuroimage 99: 103–110.

14. Fogel SM, Smith CT (2011): The function of the sleep spindle: A physiological index of intelligence and a mechanism for sleep-dependent memory consolidation. Neurosci Biobehav Rev 35: 1154–1165.

15. Mednick SC, McDevitt EA, Walsh JK, Wamsley E, Paulus M, Kanady JC, Drummond SPA (2013): The critical role of sleep spindles in hippocampal-dependent memory: A pharmacology study. J Neurosci 33: 4494–4504.

16. Kaestner EJ, Wixted JT, Mednick SC (2013): Pharmacologically Increasing Sleep Spindles Enhances Recognition for Negative and High-arousal Memories, 2013/06/19. J Cogn Neurosci. https://doi.org/10.1162/jocn_a_00433

17. Boer M De, Nijdam MJ, Jongedijk RA, Bangel KA, Olff M, Hofman WF, Talamini LM (2019): The spectral fingerprint of sleep problems in post-traumatic stress disorder. 1–9.

18. del Giudice R, Lechinger J, Wislowska M, Heib DPJ, Hoedlmoser K, Schabus M (2014): Oscillatory brain responses to own names uttered by unfamiliar and familiar voices. Brain Res 1591: 63–73.

19. Warby SC, Wendt SL, Welinder P, Munk EGS, Carrillo O, Sorensen HBD, et al. (2014): Sleep-spindle detection: Crowdsourcing and evaluating performance of experts, non-experts and automated methods. Nat Methods 11: 385–392.

20. American Psychiatric Association (1994): Diagnostic and Statistical Manual of Mental Disorders (IV).

21. Iber C (n.d.): The AASM Manual for the Scoring of Sleep and Associated Events: Rules, Terminology and Technical Specifications. Am Acad Sleep Med.

22. Schabus M, Gruber G, Schabus M, Gruber G, Parapatics S, Sauter C, et al. (2005): Schabus, M. et al. Sleep spindles and their significance for declarative memory Sleep Spindles and Their Significance for Declarative Memory Consolidation. 1479–1485.

23. Fernandez LMJ, Lüthi A (2020): Sleep Spindles: Mechanisms and Functions. Physiol Rev 100: 805–868.

24. Mednick SC, McDevitt EA, Walsh JK, Wamsley E, Paulus M, Kanady JC, Drummond SP (2013): The critical role of sleep spindles in hippocampal-dependent memory: a pharmacology study, 2013/03/08. J Neurosci 33: 4494–4504.

25. Helm E Van Der, Gujar N, Nishida M, Walker MP (2011): Sleep-Dependent Facilitation of Episodic Memory Details. 6. https://doi.org/10.1371/journal.pone.0027421

26. Rauchs G, Schabus M, Parapatics S, Bertran F, Clochon P, Hot P, et al. (2008): Is there a link between sleep changes and memory in Alzheimer’s disease? Neuroreport 19: 1159–1162.

27. Piantoni G, Halgren E, Cash SS (2017): Spatiotemporal characteristics of sleep spindles depend on cortical location. Neuroimage 146: 236–245.

28. Cox R, Hofman WF, de Boer M, Talamini LM (2014): Local sleep spindle modulations in relation to specific memory cues. Neuroimage 99: 103–110.

29. Mölle M, Born J (2011): Slow oscillations orchestrating fast oscillations and memory consolidation. 193: 93–110.

30. Cox R, Hofman WF, Talamini LM (2012): Involvement of spindles in memory consolidation is slow wave sleep-specific, 2012/06/16. Learn Mem 19: 264–267.

31. Cox R, van Driel J, de Boer M, Talamini LM (n.d.): Slow oscillations during sleep coordinate interregional communication in cortical networks. J Neurosci.

32. Besedovsky L, Ngo HV V., Dimitrov S, Gassenmaier C, Lehmann R, Born J (2017): Auditory closed-loop stimulation of EEG slow oscillations strengthens sleep and signs of its immune-supportive function. Nat Commun 8. https://doi.org/10.1038/s41467-017-02170-3

33. Mader EC, Mader ACL (2016): Sleep as spatiotemporal integration of biological processes that evolved to periodically reinforce neurodynamic and metabolic homeostasis: The 2m3d paradigm of sleep. J Neurol Sci 367: 63–80.

34. Tempesta D, Socci V, De Gennaro L, Ferrara M (2018): Sleep and emotional processing. Sleep Med Rev 40: 183–195.

35. Talamini LM, Bringmann LF, de Boer M, Hofman WF (2013): Sleeping Worries Away or Worrying Away Sleep? Physiological Evidence on Sleep-Emotion Interactions. PLoS One 8: 62480.

36. Jones S, Castelnovo A, Riedner B, Flaherty B, Prehn-Kristensen A, Benca R, et al. (2021): Sleep and emotion processing in paediatric posttraumatic stress disorder: A pilot investigation. J Sleep Res 1–11.

37. Van Liempt S, Arends J, Cluitmans PJM, Westenberg HGM, Kahn RS, Vermetten E (2013): Sympathetic activity and hypothalamo-pituitary-adrenal axis activity during sleep in post-traumatic stress disorder: A study assessing polysomnography with simultaneous blood sampling. Psychoneuroendocrinology 38: 155–165.

