## Supplemental information for "Sleep spindle dynamics suggest over-consolidation in post-traumatic stress disorder"

S1
To examine whether spindle parameters of PTSD patients versus trauma exposed controls were similarly distributed, the density distribution of each spindle parameter (duration, peak amplitude, wax amplitude, wane amplitude) was compared between the PTSD patient and control sample, using a two-sample Kolmogorov-Smirnov test. No differences were found in the shape of the distributions (p>0.1).

**A	B**

**C	D**

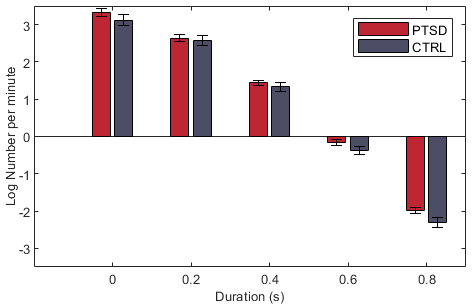

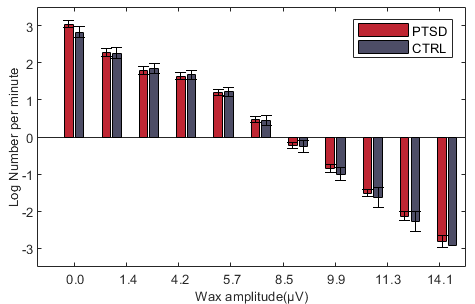

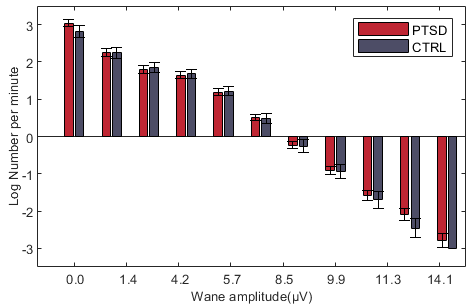

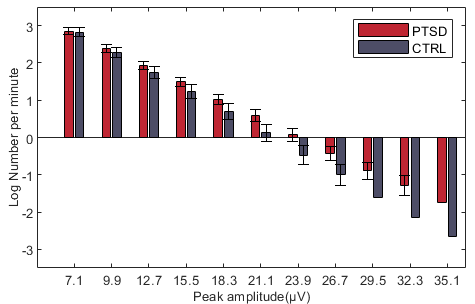

**Figure S1** Spindle event density distributions of PTSD patients versus controls, displaying the number of spindle events per minute for peak amplitude (A), duration (B), wax amplitude (C), wane amplitude (D). Logarithmic transformations were performed for visual clarity, statistics were, however, performed on the raw, nonlogarithmic data.
